# Adaptive efficient coding of correlated acoustic properties

**DOI:** 10.1101/548156

**Authors:** Kai Lu, Wanyi Liu, Kelsey Dutta, Jonathan B. Fritz, Shihab A. Shamma

## Abstract

Natural sounds such as vocalizations often have co-varying acoustic attributes where one acoustic feature can be predicted from another, resulting in redundancy in neural coding. It has been proposed that sensory systems are able to detect such covariation and adapt to reduce redundancy, leading to more efficient neural coding. Results of recent psychoacoustic studies suggest that, following passive exposure to sounds in which temporal and spectral attributes covaried in a correlated fashion, the auditory system adapts to efficiently encode the two co-varying dimensions as a single dimension, at the cost of lost sensitivity to the orthogonal dimension. Here we explore the neural basis of this psychophysical phenomenon by recording single-unit responses from primary auditory cortex (A1) in awake ferrets exposed passively to stimuli with two correlated attributes in the temporal and spectral domain similar to that utilized in the psychoacoustic experiments. We found that: (1) the signal-to-noise (SNR) ratio of spike rate coding of cortical responses driven by sounds with correlated attributes was reduced along the orthogonal dimension; while the SNR ratio remained intact along the exposure dimension; (2) Mutual information of spike temporal coding increased only along the exposure dimension; (3) correlation between neurons tuned to the two covarying attributes decreased after exposure; (4) these exposure effects still occurred if sounds were correlated along two acoustic dimensions, but varied randomly along a third dimension. These neurophysiological results are consistent with the Efficient Learning Hypothesis and may deepen our understanding of how the auditory system represents acoustic regularities and covariance.

**Significance:** In the Efficient Coding (EC) hypothesis, proposed by Barlow in 1961, the neural code in sensory systems efficiently encodes natural stimuli by minimizing the number of spikes to transmit a sensory signal. Results of recent psychoacoustic studies are consistent with the EC hypothesis, showing that following passive exposure to stimuli with correlated attributes, the auditory system adapts so as to more efficiently encode the two co-varying dimensions as a single dimension. In the current neurophysiological experiments, using a similar stimulus design and experimental paradigm to the psychoacoustic studies of Stilp and colleagues (2010, 2011, 2012, 2016), we recorded responses from single neurons in the auditory cortex of the awake ferret, showing adaptive efficient neural coding of correlated acoustic properties.

## Introduction

In the Efficient Coding (EC) Hypothesis, Barlow conjectured that spikes in sensory systems form an efficient code to represent natural stimuli and that sensory processing is optimized for natural stimuli (Barlow, 1961). Consistent with EC, there is evidence that neurons in the auditory and visual system are indeed developmentally and evolutionarily well adapted, if not optimized, for encoding sounds and images found in nature (Olshausen and Field, 1997; Vinje and Gallant, 2000; Lewicki, 2002; Smith and Lewicki, 2006). However, since many natural sounds and images have correlated attributes, hence one stimulus feature can often be predicted from another, resulting in coding redundancy. According to EC, efficient sensory systems should remove or reduce redundancies in the sensory input and in order to do so, it has been proposed that sensory systems are able to actively recalibrate coding in order to enhance coding efficiency (Barlow and Földiák, 1989) and some recent experiments in the visual system support this dynamic version of EC (Coen-Cagli et al 2015).

Additional recent support for the active EC hypothesis in the auditory system comes from psychoacoustic studies (Stilp et al., 2010, Stilp and Kluender, 2011, 2012, 2016) in which subjects either received passive exposure to, or provided continuous discrimination judgments for sets of complex sounds with covarying amplitude attack-decay ratios and spectral-shapes. Following passive exposure, and also over the course of active discrimination, subjects’ acuity in discriminating of sounds along the correlated dimensions remained intact, but discrimination of pairs on the orthogonal dimension was significantly impaired when those pairs were proximal to the principle vector of covariance. These findings suggested an interpretation whereby experience with correlated attributes (attack-decay and spectral shape) induced the auditory system to collapse the covarying dimensions into a single dimension at the expense of lost sensitivity to the orthogonal dimension. In the present study, we test whether the neural correlates of these experiments replicate this phenomenon. We do so by measuring neural responses in primary auditory cortex (A1) of awake ferrets, employing the same passive stimulus exposure paradigm as used in the original study of Stilp and colleagues (Stilp, et al. 2010).

The current study comprises three key experiments. In <u>Experiment 1</u>, we measured the baseline auditory cortical responses to sounds with a correlation between two acoustic attributes. We found that cortical responses became adapted to sounds along the dimension of the correlated attributes. And the signal-to-noise ratios along this dimension remained intact, while those along the dimension *orthogonal* to it became *depressed*. Then, consistent with predictions of efficient coding, the mutual information of spike temporal coding in auditory neurons *increased* along the exposure dimension after exposure. Finally, correlation between neurons tuned to the same two attributes *decreased* after exposure. <u>Experiment 2</u> tested whether passive exposure to stimuli varying along a single dimension, i.e., holding other parameters constant, induced effects similar to those for the correlation dimension in Experiment 1, testing whether the effects found in Experiment 1 were due to simple adaptation or whether they required a covariance between two attributes. Finally, <u>Experiment 3</u> tested whether the exposure effects shown in Experiment 1 still persisted if exposure sounds were correlated between two attributes, while a third attribute varied along a separate additional dimension (e.g., the fundamental frequency).

## Method

### Subjects

Experiments used adult female ferrets (N = 4) housed on a 12:12h light-dark cycle. Ferrets used in this study were previously trained on unrelated auditory tasks (Lu et al., 2017). Neurophysiological recording sessions (4-8 hours) occurred on two non-consecutive days per week. Ferrets were typically recorded from during one daily session for 2d per week and obtained ad libitum water on other days. All procedures were in accord with National Institutes of Health policy on experimental animal care and use and conformed to a protocol approved by the Institutional Animal Care and Use Committee (IACUC) of University of Maryland.

### Surgeries

In order to stabilize the head for electrophysiological recording, a headpost was implanted in a surgery that occurred at least one month before the initiation of recordings. Animals were anesthetized with isoflurane (2% in oxygen), and a customized stainless steel head-post was surgically implanted on the skull under aseptic conditions. The skull over the auditory cortex was exposed and covered with a thin layer of Heraeus Kulzer Charisma (1 mm) surrounded by a thicker wall built with UV-curable Charisma (3 mm thick, 10 mm diameter). After recovery from surgery, animals were habituated to a customized head-fixed holder. Two days before electrophysiological recording, a small craniotomy (1-2 mm diameter) was made above primary auditory cortex. At the beginning and end of each recording session, the craniotomy was thoroughly rinsed with sterile saline. At the end of a recording session, the craniotomy and well was filled with topical antibiotics that were rotated on a weekly basis (Baytril and Cefazolin). The area containing the hole was then filled with sterile vinyl polysiloxane impression material (Examix NDS) that maintained a tight seal between experiments. After recordings were completed in the original craniotomy, adjacent 0.5 mm bone sections were removed over successive months of neurophysiological recording so that eventually the enlarged craniotomy (~4 mm diameter) encompassed the entire primary auditory cortex.

### Electrophyiological recording

Electrophysiological recordings were made in a double-walled soundproof room (IAC, New York). The awake animal was placed in a horizontal, lexan tube, and the implanted headpost was used to stabilize and fix the head in position, relative to a stereotaxic frame. Recordings were conducted in primary auditory cortex (A1) of both left and right hemispheres over a period of 3-6 months. For each recording, 4-8 tungsten microelectrodes (2–3 M MOhms, FHC) were introduced through the craniotomies and controlled by independently moveable drives (Electrode Positioning System, Alpha-Omega). Raw neural activity traces were amplified, filtered, and digitally acquired by a data acquisition system (AlphaLab, Alpha-Omega). Multiunit neuronal activity (including all spikes that rose above a threshold level of 3.5 SDs of baseline noise) was monitored online. In addition, single units were isolated online to monitor single neuron responses. Bandpass noise (0.2 s, 1 octave) and pure tone (0.2 s duration) stimuli were presented to search for responsive sites. Because of evidence that neurons in supragranular layers (II-III) of A1 show greater plasticity than neurons in deeper layers (Francis et al., 2018), most of the recording depths in this study were within 100-400 microns of the cortical surface, and electrodes were advanced only to the most superficial position where single-unit responses to bandpass noise and tones were found. Once clear, stable auditory responses were obtained, the experimental stimuli were presented. All stimulus amplitudes were presented at 65 dB SPL from a speaker placed 1 meter in front of the animal. After recordings were completed, single units were isolated again by off-line customized spike-sorting software, which was based on a PCA and template-matching algorithm (Meska-PCA, NSL).

### Auditory Stimuli and Experimental design

Experiments were organized into three tracks. In each, animals were exposed to a particular set of acoustic stimuli that differed in fundamental ways. In Experiment 1, two acoustic attributes were correlated and we explored the effects of this correlation on the coding of the stimuli. Experiment 2 was a control experiment where only one attribute was varied in order to test whether simple adaptation of the responses could explain results in Experiment 2. In Experiment 3, the same acoustic attributes used in Experiment 1 were correlated, but now we explored the effects (or lack thereof) of having in addition a third independently varying (uncorrelated) attribute.

## Experiment 1

### Stimuli

In each recording session, we generated a stimulus matrix with two manipulated attributes: amplitude modulation rate (AM) and peak frequency of the spectral envelope (SP). All stimuli were harmonic complexes with an initially flat spectrum modulated/filtered by the two attributes. Exposure stimuli were generated with orthogonal correlations between attributes. AM and SP were either positively correlated (increasing rate with increasing peak frequency (**Figure 1A**, represented by red dots in **Figure 1C**) or negatively-correlated (increasing rate with decreasing peak frequency, blue dots in **Figure 1C**). “Test” stimuli were generated with all combinations of the two attributes, so that they were uniformly distributed in the 2-dimensional space of the two parameters (represented by the black dots in **Figure 1C**).

**Figure 1.**
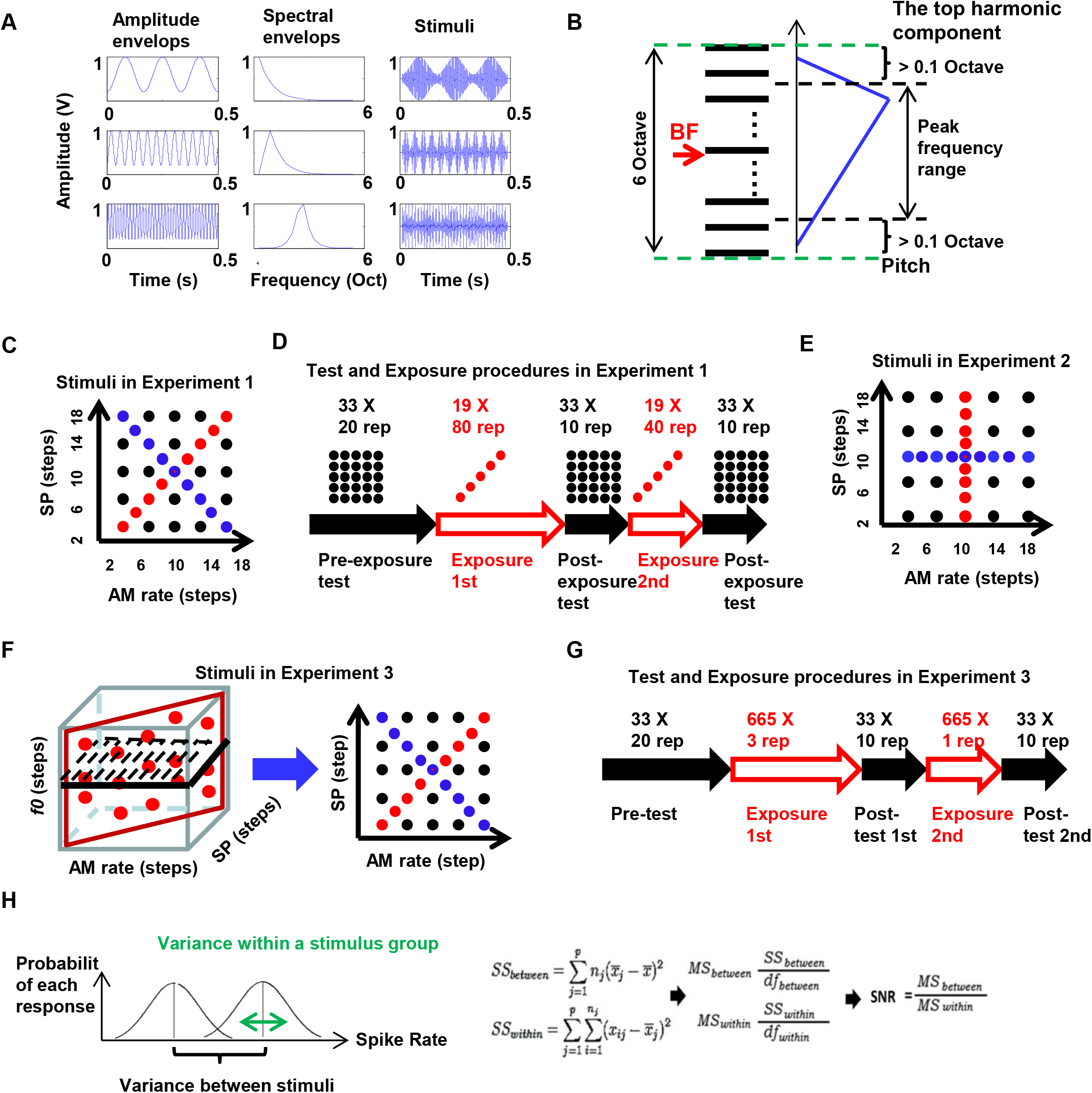
A. Schematic illustration of how stimuli were generated. Amplitude envelopes with different AM modulation rates were generated (left column). Spectral envelopes with different peak frequencies were generated (middle column). Sounds were generated from the two properties (right column). B. Illustration of how stimuli parameter was chosen. Black horizontal lines indicate harmonic components consisting of the basic structure of a stimulus. The center of the harmonic tone was chosen around the BF of recorded neurons, indicated by the red arrow. The blue triangle represented one of the 19 spectral envelopes used to generate stimuli. The limit of the peak frequencies was indicated by the two horizontal black dashed lines. The range of the harmonic structure was 6 octaves. The limit of the lowest harmonic component (*f*0) and highest component, were indicated by the green dashed line, at least 0.1 octave away from the limits of the peak frequencies. C. Testing stimuli space in Experiment 1. Notice that, in half of recordings, 19 exposure stimuli had positive correlated attributes and thus were lined up with those red dots. In this case, the orthogonal dimension was indicated by blue dots. In the other half of recordings, the opposite was true: the exposure dimension was lined up with blue dots and orthogonal dimension was lined up with red dots. Two type of exposure were shuffled across recordings in a balanced design. D. Exposure and testing procedure in Experiment 1. Black arrows indicated the testing session. Red arrows indicated the exposure session. The matrix of black dots and red dots on the top of errors illustrated stimuli presented in each session. E. Testing stimuli space in Experiment 2. In half of recordings, sounds with constant AM and varying SP (lined up with red dots) were exposure sounds, while those with constant SP and varying AM (lined up with blue dots) were on the orthogonal dimension. In the other half of recordings, opposite was true. Two type of exposure were also shuffled in a balanced design. F. Testing stimuli space in Experiment 3. The red parallelogram indicated exposure sounds, which had perfect positive or negative correlation between AM and SP, and varied *f*0. The black parallelogram with dashed lines indicates testing sounds, in which only one *f*0 was randomly chosen for testing in each recording. The structure of the testing space is same as in Experiment 1, indicated on the right by the blue arrow. G. Exposure and testing procedure in Experiment 3. Black arrows indicated the testing session. Red arrows indicated the exposure session. The total stimuli number in exposure was approximately the same as that in Experiment 1 (2280 in Experiment 1; 2660 in Experiment 3). H. Illustration of how SNR of spike rate coding for the exposure dimension and orthogonal dimension was calculated. The bell-shaped curves illustrate the possible spike rate variation within the group, the sum of which across all groups yielded the “SS within”. The distance between the centers of bell curves indicated the variance between the groups, explained by stimulus differences along the given dimension. The sum of all such variance yielded the “SS between”. Then SNR was calculated from “SS within” and “SS between” as shown in the equation on the right.

At the beginning of each recording session, several neurons were isolated; best frequencies (BFs) measured; and, median BFs estimated. Once the frequency tuning was measured, tone-complexes ranging across +/− 3 octaves around median BF (**Figure 1B**) were created. Nineteen frequencies were defined as SP peaks with equal (1/3 octave) steps across this range, and 19 triangular SP filter shapes were designed (three examples are shown in the second column of **Figure 1A**). Nineteen AM modulation rates were set at equal log steps from 5 Hz to 120 Hz (depicted in the first column of **Figure 1A**). In each recording session, the fundamental frequency (*f*0) of the harmonic complex was randomly selected from a range of 200 to 500 Hz following the criterion that *f*0 must be 0.1 octave lower than the lowest peak frequency of spectral functions (green dashed line on the bottom of **Figure 1B**). Roving *f*0 across sessions minimized effects of frequency-specific influences (e.g., peak harmonic relative to peak of spectral envelope) and memory. Stimuli were 500-ms duration with 5-ms cosine onset and offset ramps, and sampled at 40 kHz. Test stimuli included 25 stimuli selected from the combination of the steps 2, 6, 10, 14, 18 from both two dimensions forming a 5 X 5 matrix uniformly sampled from the 19 X 19 exposure matrix. Finally, because all exposure stimuli were located along diagonals of the stimulus matrix, additional 8 *test stimuli* were selected along these two diagonals of the stimuli matrix at steps 4, 8, 12, 16.

### Passive exposure and testing procedure

In Experiment 1, two ferrets were first tested with the full stimuli matrix shown in **Figure 1C** to measure neurons’ tuning properties before exposure. This was followed by exposure to repetitions of stimuli along one diagonal (red dots or blue dots). next, a post-exposure test employed the full matrix as in the Pre-exposure test to measure effects of passive exposure to stimuli with correlated attributes. Because of uncertainty regarding the duration of persistence of exposure effects, we repeated the Exposure/Post-exposure sequence. Exact procedures for stimulus presentations are summarized in **Figure 1D**: (1) Pre-exposure test stimuli, including the 33 testing stimuli ((5×5) + 8) presented 20 times each with 1s silence (ISI) between sounds – all over a period of 16.5 min; (2) First passive exposure session: 19 exposure stimuli presented 80 times each with 0.25s ISI over a period of 19 min. (3) Post-exposure test using the same stimuli of the Pre-exposure test with 10 repetitions over a period of 8.25 min. (4) Second exposure session was conducted immediately afterwards using the same stimuli repeated 40 times to reassess the effects of the first exposure over a period of 9.5 min. (5) Post-exposure test stimuli repeated 10 times over a period of 8.25 min. Data collected from the two Post-exposure tests (3,5) were combined for comparison with the Pre-exposure test (1).

### Experiment 2

A possible confound is that the effects observed in Experiment 1 could simply have been due to adaptation to repetitive sounds. As a control, two more ferrets were tested (Experiment 2) with 19 sounds during the Passive exposure phase, which varied along a *single* dimension (AM or SP, balanced across recordings). The parameter of the other dimension (orthogonal dimension) was held constant in a given recording session (**Figure 1E**) but was different across sessions. Testing stimuli In Experiment 2 included a 5 X 5 stimuli matrix as in Experiment 1. In contrast with Experiment 1, however, in Experiment 2 the additional 8 testing stimuli were sampled at a single row or column of the stimuli (rather than at the diagonals) at the step 4, 8, 12, 16. The exposure, test procedures and ISI were the same as in Experiment 1, except that exposure stimuli were along the vertical or horizontal direction, rather than a diagonal direction. By comparing the results of Experiments 1 and 2, we could assess the specific effects of the two correlated attributes.

### Experiment 3

In Experiment 3, we tested whether the exposure effects caused by covariation in two acoustic dimensions was sustained in the presence of substantial variation in a third acoustic dimension that introduced widely varying physical acoustic properties. Stilp and Kluender (2012) interpreted such a finding as evidence for increasingly non-isomorphic representation of stimulus redundancy. Here, we introduced this variation by varying *f*0 from trial to trial. Two ferrets were tested in Experiment 3. As in Experiment 1, 19 AM and SP combinations were selected, in which the two dimensions were either positively or negatively correlated. In addition, 35 *f*0 parameters were generated in 2% increments between the two nearest steps. Fundamental frequency range was restricted by the criteria that the lowest *f*0 had to be 1.5 octave higher than the maximum AM modulation rate, and the highest *f*0 had to be 0.1 octave lower than the lowest spectral peak selected for the SP dimension. A matrix of 665 (19 X 35) stimuli with 19 AM/SP combinations as one dimension and 35 *f*0s as the second dimension were created as exposure stimuli (red vertical rectangular in **Figure 1F**). For testing, we randomly selected one *f*0 out of the 35 *f*0 values in each recording. Then, testing stimuli were created based on the selected *f*0 in the exactly same way as in Experiment 1 (Black horizontal rectangular in **Figure 1F**). Thus, the *exposure* stimuli in Experiment 3 existed in a two dimension space that was orthogonal to the space of *testing* stimuli. Pre-exposure and Post-exposure test procedures were the same as in Experiment 1. Note that in Experiment 1, all 19 AM/SP combinations in passive exposure were presented 120 times. In order to ensure approximately equal covariance exposure, all 665 exposure stimuli in the first passive exposure session of Experiment 3 were presented three times (**Figure 1G**). Then 665 stimuli were presented one more time in a second passive exposure session (totally 140 repeats of 19 AM/SP combinations, but with different *f*0s).

### Data analysis

As indicated above, we isolated single units off-line from our recordings, and all the data analysis was conducted on isolated single neurons from A1. For each trial, we measured spike rate in a 650-ms response window (50 after stimulus *onset* – 200 ms after stimulus *offset*) and the spike rate in a baseline window (200 ms *before* stimulus onset). The response amplitude was defined as the spike rate in the response window minus the baseline spike rate, and was averaged over 20 repetitions. Any trials with response amplitude less than or greater than 5 standard deviations from mean amplitude were excluded from analysis. Response amplitudes in the Pre-exposure test and Post-exposure test were calculated separately. Trials from two Post-exposure tests (3,5) were combined for analysis. Baseline spike rates were compared across all three test sessions. Units with significant changes between baseline measures were excluded from analysis.

#### Quantification of signal-to-noise ratio in spike rate coding

We employed three response measures and explored how each of these measures of neural activity was or was not affected by exposures in each of the three experiments.

The first measure is signal-to-noise ratio (SNR) of spike rate coding, defined as the ratio of the response variance to stimuli along one dimension, to the overall variance within each stimulus group (**Figure 1H**). This analysis was done for each neuron. The calculation procedure resembled a two-way ANOVA, in which stimulus levels along the exposure (correlated) dimension and the orthogonal (uncorrelated) dimension were treated as two independent variables, and the response amplitude of each stimulus was treated as the dependent variable.

The same analysis was conducted for data from the Pre-exposure test and the Post-exposure tests, and differences in SNRs (change in SNRs) before and after exposure were measured. SNR changes due to exposure were analyzed separately for exposure dimension and for orthogonal dimension. In addition, SNRs themselves were compared between the Pre-exposure test and the Post-exposure test. Because the distribution of SNRs did not fully satisfy criteria for parametric tests, Wilcoxon test (a non-parametric version of paired t-test) was performed to compare SNRs before and after exposure.

The second response measure is mutual information (MI) carried by spike trains within the response window prior to and following exposure. MI was defined as the difference between the entropy associated with *all* responses from a neuron vs the entropy of responses given a stimulus. First, spike trains were segmented with a series of 100 ms bins, in which occurrence of spikes in a bin was indicated as 1 and absence as 0. In this way, the spike train in each trial (600 ms duration) was converted into a binary word with length of 6 (bins beginning 50ms after stimulus onset and continuing until 150 ms after stimulus offset – stimuli were 500 ms duration). Then, the entropy of spike trains were calculated from the distribution of *all* binary words in response to *all* stimuli, as

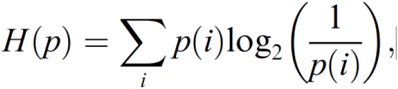

In which, p is the probability of observing one binary word (representing a specific spike train pattern) from *all* responses recorded from a given neuron. Then, the same calculation was done only for responses to a given stimulus, the sum of which yield the entropy of joint distribution between stimulus and response. MI was the difference between the two entropy values. We followed standard procedures described in Nelken et al, 2007: MI was calculated from randomly selected subsets of data of four sizes (40%, 60%, 80% and 100%). Then a linear regression between MIs and the reciprocal of the sample size in each data set was calculated. The intercept of the linear regression provided an estimate of the unbiased MI. MI was calculated for both the exposure dimension and orthogonal dimension in each neuron. Analyses were performed in the same way for each neuron before and after exposure. Wilcoxon tests and binomial tests were used to compare MI obtained before and after exposure.

Finally, the third response measure was the correlation coefficient between the tuning functions to the AM and SP parameters for the same neuron or neuron pairs recorded simultaneously in each recording (neuron pairs were distinct neurons recorded simultaneously either from adjacent electrodes or separate single units (based on waveform and tuning) recorded from the same electrode). For analysis of correlation in the same neuron, the tuning functions to AM and SP were calculated based on the averaged response at each AM/SP level. The correlation between the two functions was then calculated (with Spearman correlation) for each neuron, and for each Pre-exposure and Post-exposure test. Next, we computed the differences in correlation coefficients obtained from all neurons before and after exposure. Finally, correlation coefficients from recordings with different stimulus exposures – e.g., positively versus negatively correlated – were computed and compared with each other. Because the population of correlation coefficients was not normally distributed, the Mann–Whitney U test (non-parametric version of two-sample t-test), which provides a more conservative evaluation than a traditional t-test, was used for the positive/negative comparison. To analyze correlations between AM and SP from different simultaneously recorded neurons, we first paired neurons recorded simultaneously in each session, one for AM and one for SP. Then, the correlation between AM and SP functions was calculated for the neuron pairs and averaged across all pairs. The differences between correlation coefficients before and after exposure were compared between recordings with exposure to positively and negatively correlated stimuli.

## Results

### Experiment 1: effects of exposure to stimuli with two correlated attributes

We first examined the effects of exposure on the neural responses to stimuli with two correlated attributes. We first consider (a) changes in the tuning properties of the neurons with respect to the AM and SP parameters, then (b) examine changes in coding quality (or SNR), followed by (c) changes in MI, and finally describe (d) the changes in inter-neuronal correlations among simultaneously recorded neurons.

#### a. Effect of exposure on tuning properties

In Experiment 1, we recorded from 65 single-units in primary auditory cortex (A1) of two ferrets. The differences between overall responsiveness of neurons after exposure to all stimuli were not significantly different (Wilcoxon test: *z* = −0.8332. *p* = 0.404). However, there was a significant change when we contrasted two groups of stimuli separately. Specifically, contrasting responses to “exposure” stimuli (combinations within the green rectangle in **Figure 2A**) versus the non-exposure stimuli (combinations within blue triangles in **Figure 2A**), revealed a clear effect of adaptation: responses to the exposure stimuli (both positively or negatively correlated) significantly *decreased* in the Post-exposure tests (Wilcoxon test: *z* = −2.6. *p* = 0.001, left-histogram in **Figure 2B**, and left-box in **Figure 2C**), while responses to non-exposure stimuli significantly *increased* after exposure (Wilcoxon test: *z* = −2.4. *p* = 0.016, right-histogram in **Figure 2B**, the right-box in **Figure 2C**).

**Figure 2.**
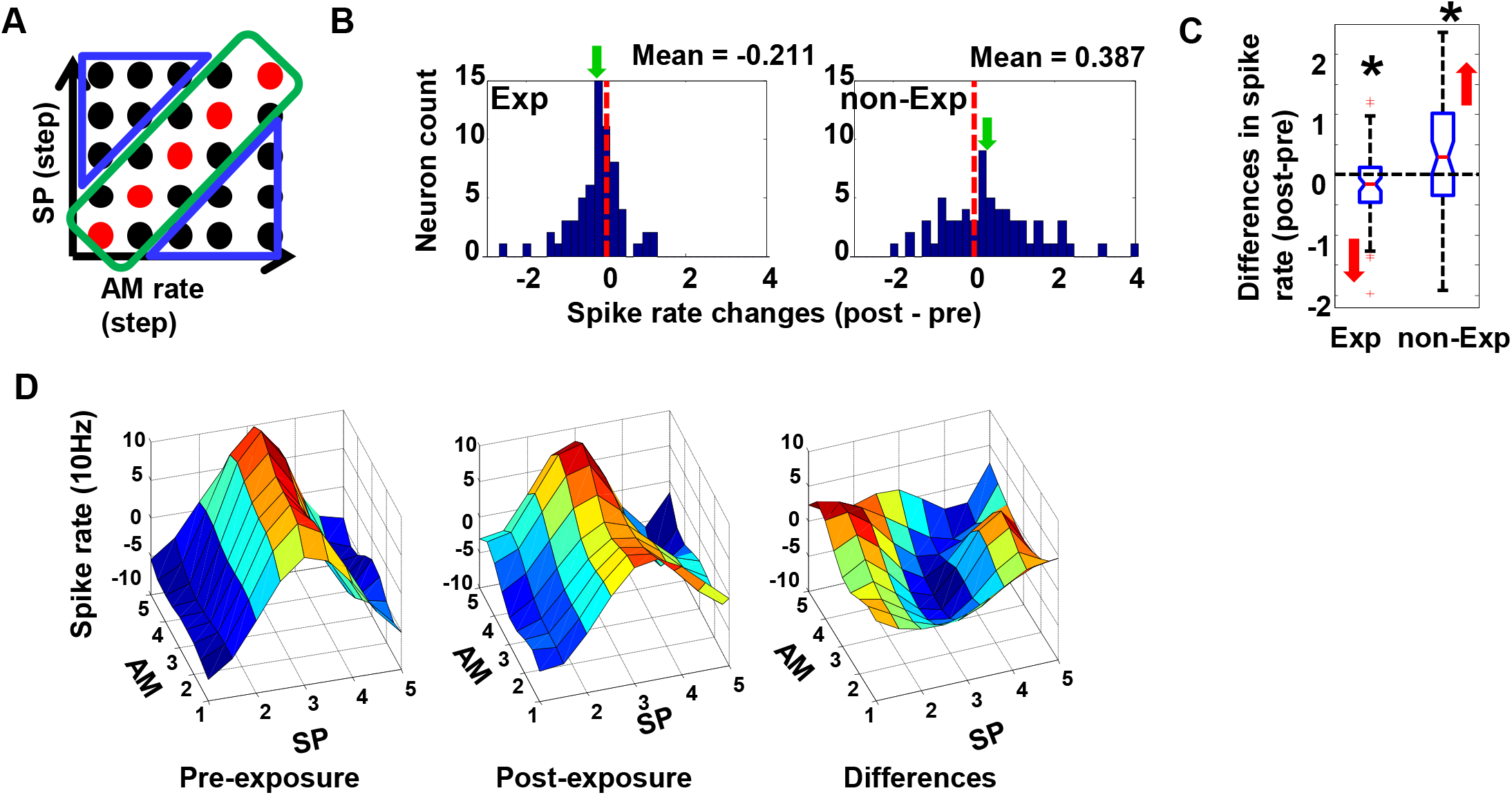
A. Exemplars of stimuli used in tuning analysis in Experiment 1. Green rectangular indicated exposure stimuli or nearby, from which the spike rates were averaged and compared between tests before and after exposure. Two blue triangular indicated non-exposure stimuli, (far from exposure stimuli), from which the spike rates were averaged for comparison. B. Histogram of spike rate changes after exposure in Experiment 1 (exposure stimuli and nearby on the left and non-exposed on the right). The red dashed line indicates zero. The distribution of spike rate changes for exposure stimuli was significantly biased to the negative side, while the distribution for non-exposure stimuli was biased to the positive side. The green arrow indicates the mean (also indicated on the top of histogram) for each distribution. C. Box plot of spike rate changes after exposure in Experiment 1 (exposure stimuli and nearby on the left and non-exposed on the right). The red line in the middle and the notch indicated the median and its 95% confidence interval. The bottom and top edges of the box indicate the 25th and 75th percentiles. The whiskers indicate the range. Red cross indicate possible outliers. The star on the top indicates that the distribution was significantly different from zero. The red arrow on the side indicates the direction of significant changes. Spike rates significantly decreased for the exposure stimuli and significantly increased for non-exposure stimuli. D. Surface plot of the 2-dimensional tuning map in Experiment 1: before exposure (left), and after exposure (middle). The unit of AM and SP was the step used to create the stimulus matrix. The differences between the two maps were taken (post minus pre) and showed on the right. The changes of tuning map were consistent with quantification showed in Figure 2B.

This result is consistent with previous findings by Dragoi et al. (2000), where adaptation was also shown to cause lateral shifts in neuronal tuning functions, thus significantly modifying them, as illustrated by the surface plot of response amplitudes to each test stimulus in **Figure 2D**. Note that tuning map from recordings with negative correlated exposure was flipped so that the exposure dimension from two types of exposure (positive and negative) could be aligned.

#### b. Effect of exposure on the coding quality (SNR)

We also analyzed the effect of exposure on the SNR of each neuron. Early psychoacoustic studies (Stilp et al. 2010) had shown that, after passive exposure to sounds with correlated properties, the auditory system captured the covariance of the two acoustic attributes and treated them as a single perceptual dimension, while simultaneously losing discriminability along the orthogonal dimension. Therefore, we hypothesized that the (SNR) of spike rate coding along the orthogonal dimension would also be reduced after exposure.

To test this hypothesis, we calculated the SNR for each neuron, with stimulus levels along the exposure dimension and the orthogonal dimension as two independent variables (**Figure 3A**). Data from exposure to positively- and negatively-correlated properties were combined in this analysis. The results confirmed the prediction that the SNR on the exposure dimension remained unchanged (Wilcoxon test: *z* = −1.1. *p* = 0.268), while it significantly decreased along the orthogonal dimension (Wilcoxon test: *z* = −2.0. *p* = 0.042). **Figure 3B** illustrates this finding by a histogram of the normalized SNR changes (divided by the sum of the pre- and post-SNRs). On the exposure dimension (left histogram), SNR’s changes were symmetrically distributed around zero, while along the orthogonal dimension they were significantly biased to the negative side (right histogram). This pattern is also demonstrated by the box plots in **Figure 3C**. Furthermore, we did not find significant SNR changes for the interaction between the exposure and orthogonal dimensions (Wilcoxon test: *z* = −0.1. *p* = 0.909). In summary, the pattern of neuronal adaptation is consistent with the findings of the psychoacoustic studies (**Stilp et al., 2010**).

**Figure 3.**
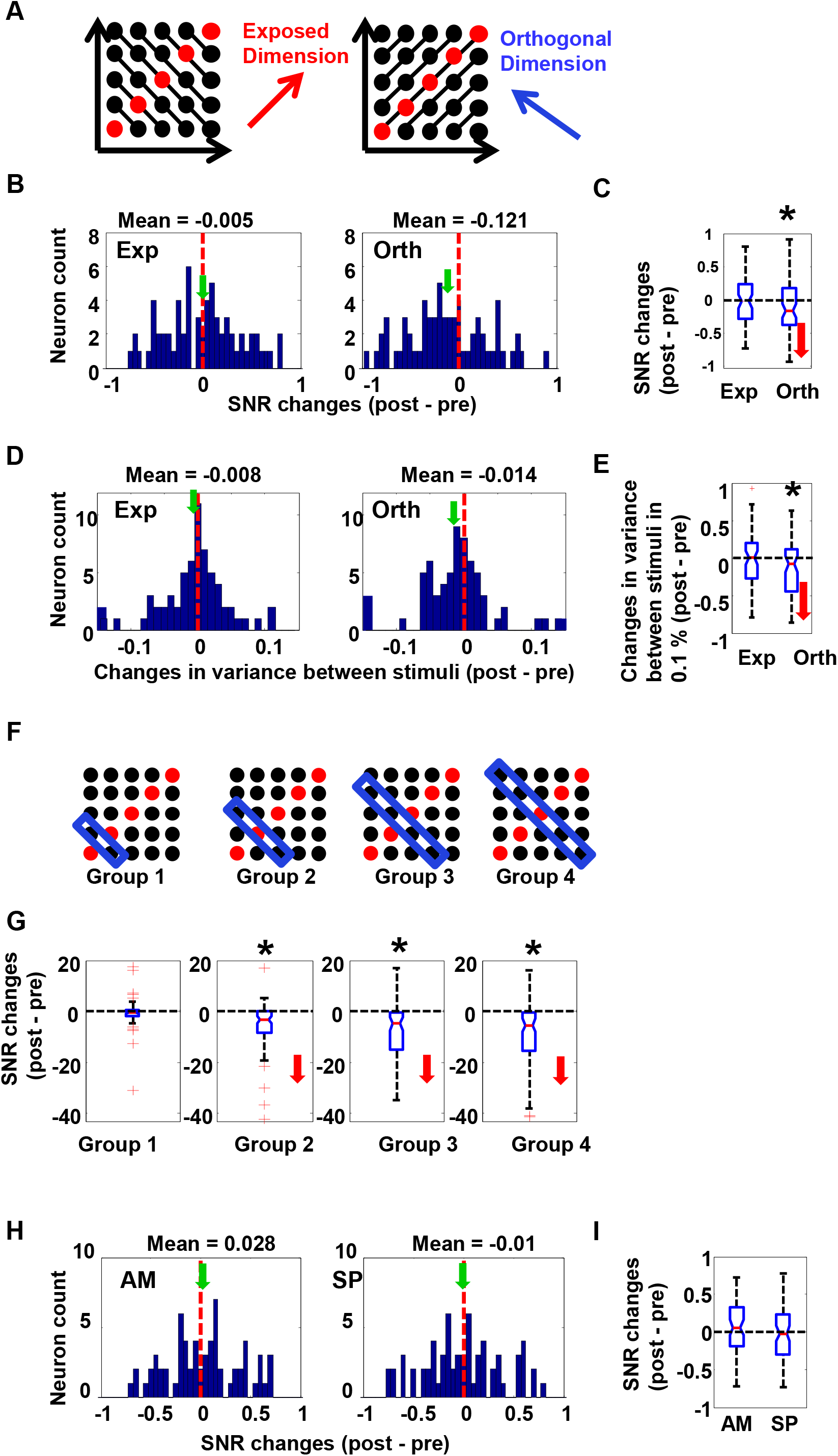
A. Schematic illustration of calculation of SNR along exposure (left) and orthogonal dimension (right). Each line connected with dots indicates stimuli to be treated as one single group (one level on the exposure or orthogonal dimension). Variance between stimuli was compared between lines (groups). The red arrow indicates the direction of the exposure dimension. The blue arrow indicates the direction of the orthogonal dimension. B. Histogram of SNR changes in Experiment 1. SNRs for exposure dimension were symmetrically distributed around zero (left, red dashed indicated zero). SNRs for orthogonal dimension were significantly reduced (right). The green arrow indicated the mean (also indicated on the top of histogram) for each distribution. The SNR change (post minus pre) was taken for each neuron and the difference was normalized by dividing by the mean of the two values. Normalization was only applied for display, while statistical analysis was done without normalization. C. Box plot of SNR changes in Experiment 1. SNRs for orthogonal dimension were significantly reduced (right). The star on the top indicated that the distribution was significantly different from zero. D. Histogram of changes in variance (in spike rates) between stimuli in Experiment 1. Only the histogram on the right (for orthogonal stimuli) was significantly biased to the negative side, resembling the plot for SNR changes (above). E. Box plot of changes in variance (in spike rates) between stimuli in Experiment 1. Variance between stimuli for orthogonal dimension was significantly reduced (right). The star on the top indicated that the distribution was significantly different from zero. F. Schematic illustration of the subgroup selection, indicated by the blue box, for the tests of SNR changes in Experiment 1. The stimulus group on the left was located at the corner of the test space, while the stimulus group on the right was located on diagonal line of the testing space. G. Box plots showed the effect of exposure on SNR changes on the orthogonal dimension in each subgroup illustrated on the top (Figure 3E). The effect was most strong when test stimuli were located on the diagonal line (Group 4, on the right), because stimuli were distributed with longest distance along the orthogonal dimension. H. Histogram of normalized SNR changes for AM (left) and SP (right) in Experiment 1. There was no significant change for either dimension. SNR changes were all symmetrically distributed around zero in both plots. The mean for each distribution is indicated both on the top of the plot and by the green arrow. I. Box plot of normalized SNR changes for AM (left) and SP (right) in Experiment 1. There was no significant change for either dimension.

Because SNR is a ratio of two variances (within/between stimuli), it was unclear what exactly caused the SNR changes, i.e., which of the two variances dominated the changes. By examining the two variances separately, we found that the SNR changes on the orthogonal dimension were due to reduced variance between its stimuli along the diagonal (**Figure 3D and Figure 3E**, Wilcoxon test: *z* = −2.1. *p* = 0.035), while variance within each stimulus remained the same after exposure (Wilcoxon test: *z* = −1.2. *p* = 0.243). Therefore, the reduction in SNR was primarily due to a reduced signal representation on the orthogonal axis (i.e., diminished response differences between stimuli), rather than changes in noise level (i.e., variance of responses within each stimulus). <u>It should be noted that t</u>he size of this SNR decrease depended on how many stimuli were distributed along the orthogonal dimension. For example, because stimuli located in the middle of the diagonal were influenced by more stimuli along the orthogonal dimension than those located near the corners (**Figure 3F**), we conjectured that stronger effects would be seen there. To test this, we calculated and compared SNRs before and after exposure for each of 4 separate groups of stimuli defined by their distance to the diagonal, and also averaged the changes from stimuli that were symmetrically placed on either side of the diagonal. As shown in **Figure 3G**, the effect of exposure along orthogonal dimension peaked at the diagonal (Wilcoxon test: *z* = −4.6. *p* = 0.001), becoming weaker towards the corners (Wilcoxon test: *z* = −1.8. *p* = 0.078). This decrease in effect size with increasing distance from the diagonal replicates psychophysical studies by Stilp and Kluender (2012, 2016). <u>Finally, we also measured the</u> SNR changes separately for stimuli along the pure AM and SP dimensions. The results revealed no significant changes along either axis (**Figure 3H and 3I**, Wilcoxon test: *z* = −0.11. *p* = 0.914 for AM, and Wilcoxon test: *z* = −1.19. *p* = 0.235 for SP), with no significant changes in interactions between them (Wilcoxon test: *z* = −0.2. *p* = 0.847).

#### c. Mutual information conveyed by spike trains increased for exposure stimuli

We next examined whether there was any gain in coding efficiency along the exposure (correlated) dimension. We did so by analyzing the temporal coding in the spike trains, as quantified by the MI between the responses and stimuli (see Methods for definition). Because test stimuli along the correlation (diagonal) axis exhibited the largest adaptive effects of exposure, we focused on them for the MI calculation (**Figure 4A**). We found that, unlike the SNR changes following exposure, the MI on the orthogonal dimension remained unchanged. As shown in histogram of **Figure 5B** (right), 49% of neurons showed increased MI vs 51% of neurons showed decreased MI after exposure on the orthogonal dimension, resulting in no significant changes (binomial test: *p* = 0.500). In contrast, as shown in the histogram of **Figure 4B** (left), 63% of neurons exhibited the increased MI on the exposed dimension. MI significantly increased on the exposure dimension (binomial test: *p* = 0.023; summary shown in the box plot in **Figure 4C**).

**Figure 4.**
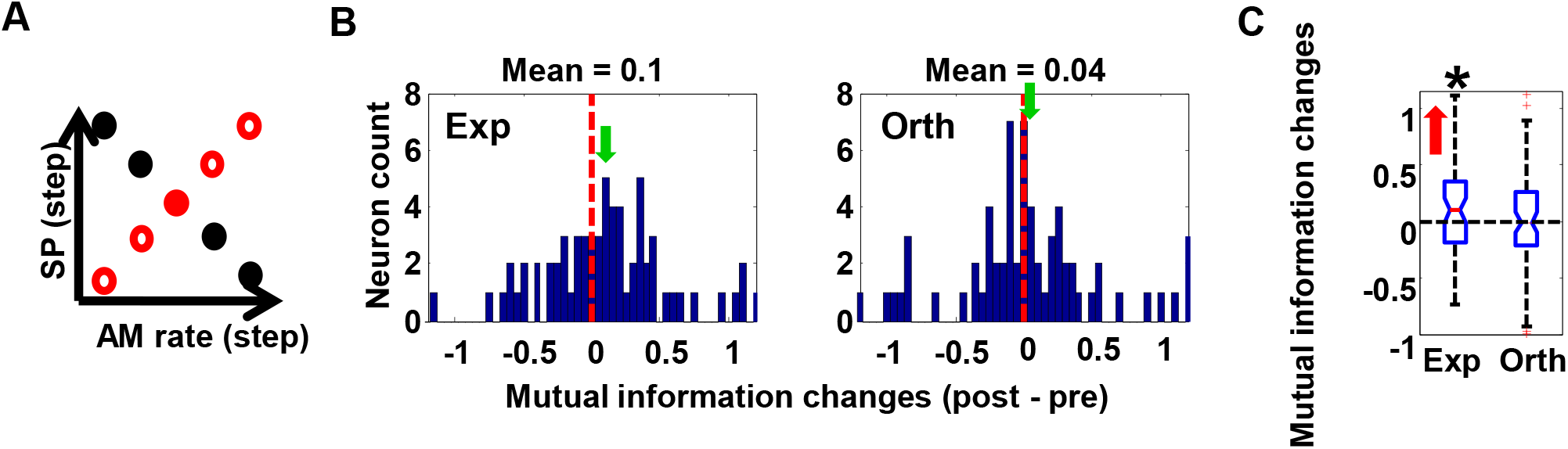
A. Schematic illustration of the stimuli tested for MI changes in Experiment 1. Red dots indicate exposure stimuli. Black dots indicate orthogonal stimuli. B. Histogram of MI changes in Experiment 1. The red dashed line indicates zero. The 63% of MIs for exposure stimuli were on the positive side (left), while the distribution of MIs for orthogonal dimension was almost symmetrically centered at zero (right). C. The box plot of MI changes in Experiment 1. MI significantly increased for exposure stimuli, showed by results of the binomial test, but not for orthogonal stimuli.

**Figure 5.**
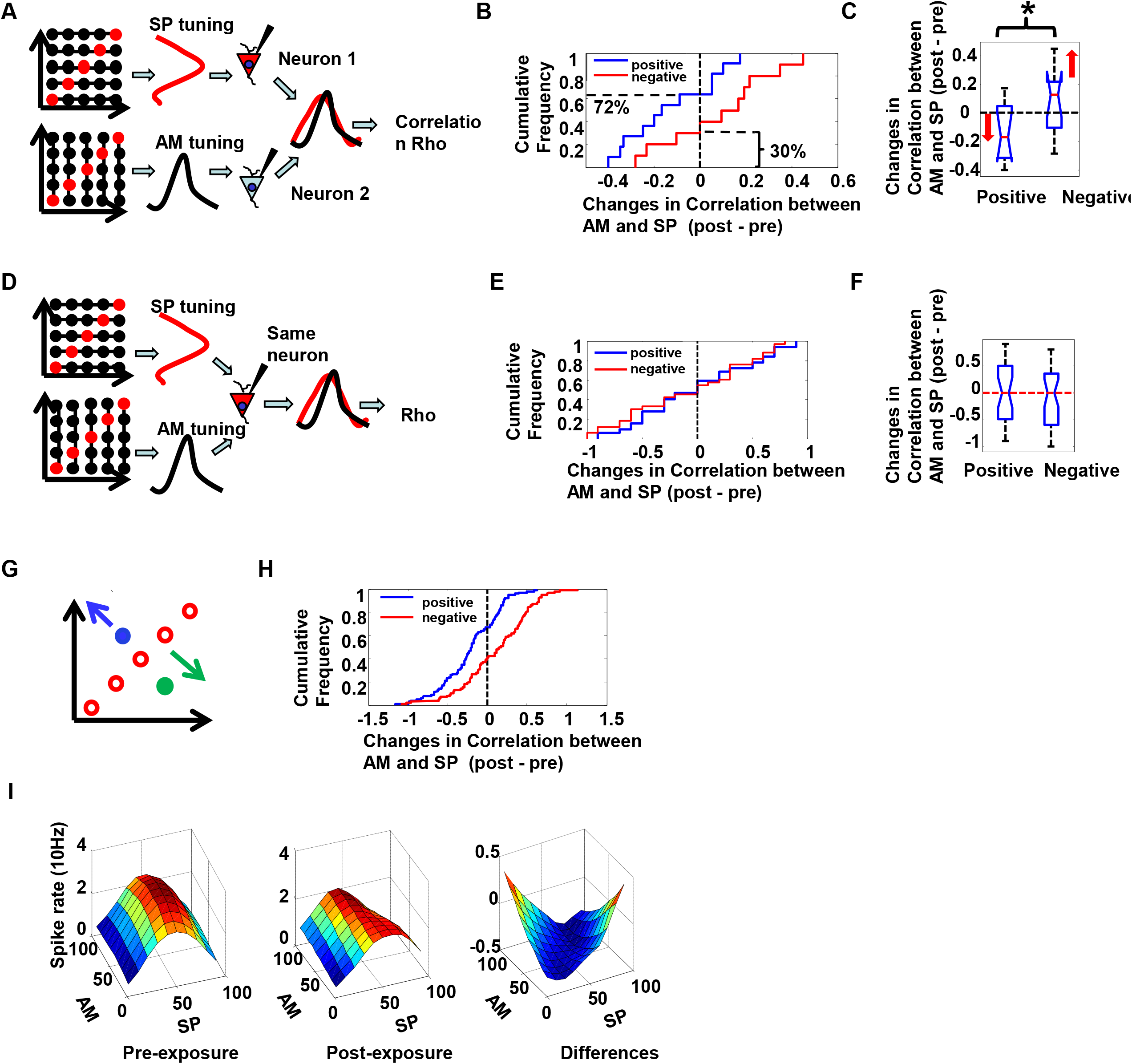
A. Schematic illustration of calculation of correlation of tuning from different neurons in Experiment 1. On the top, each line connected with dots indicated one level of SP to be averaged to create the tuning curve for SP (red bell shape curve). On the bottom, each line connected with dots indicated one level of AM to be averaged to create the tuning curve for AM (Black bell shape curve). The two tuning curves were calculated from two different simultaneously recorded neurons: Neuron 1 (red) and Neuron 2 (blue). Then, correlation was computed from the red and the blue tuning curve obtained from two neurons. At last, difference in correlation coefficient obtained before and after exposure was taken. B. Cumulative frequency distribution (left) and the box plot (right) of tuning correlation changes cross neurons in Experiment 1. The Y axis of cumulative frequency plot shows the percent of the total sample lower than (or equal to) an associate value on the X axis. The vertical black dashed line indicates zero. The blue trace shows correlation changes after exposure to positively correlated attributes. The horizontal black dashed line on the left of the blue trace indicates that correlation changes were negative in 72% (8 out of 11) recordings after exposure to positively correlated attributes. The red trace shows correlation changes after exposure to negative correlated attributes. The horizontal black dashed line on the right indicates that only the 30% (3 out of 10) of red trace was on the negative side. C. Box plot of tuning correlation changes cross neurons in Experiment 1. The bracket and the star indicates that the two distributions were significantly different from each other. D. Schematic illustration of the tuning correlation calculation for the same neuron in the Experiment 1. On the top, each line connected with dots indicated one level of SP to be averaged to create the tuning curve for SP (red bell shape curve). On the bottom, each line connected with dots indicates one level of AM to be averaged to create the tuning curve for AM (black bell-shaped curve). The two tuning curves were calculated from the same neuron. Then, the correlation was computed from the red and the blue tuning curves. E. Cumulative frequency distribution (left) and the box plot (right) of tuning correlation changes for the same neuron in Experiment 1: There was no significant change in tuning correlation either for positively correlated exposure (blue trace) or for negative correlated exposure (red trace). F. Box plot of tuning correlation changes for the same neurons in Experiment 1. G. Illustration of hypothesis that exposure to correlated attributes pushes neuron’s tuning function away from the exposure stimuli along the orthogonal direction. The blue and green dots illustrate the best tuning position of two hypothetical neurons in the stimulus matrix. The arrows indicate the direction of tuning changes. H. Results of modeling. Cumulative frequency distribution of tuning correlation changes in 200 simulated neurons. Consistent with results, correlation significantly decreased after exposure to positively correlated attributes (red trace) and significantly increased after exposure to negatively correlated attributes (blue trace), resembling results shown in Figure 5B. I. Results of modeling. Surface plot of the 2-dimensional tuning map obtained from 200 simulated neurons: before exposure (left), and after exposure (middle). The differences between the two maps were taken (post minus pre) and are shown on the right. The overall pattern of change was consistent with our results in Figure 2D.

#### d. Decorrelated tuning functions

We finally investigated exposure effects at the neuronal population level following Experiment 1 exposure paradigm. We first assume that if two neurons are tuned to different sound attributes, e.g., AM or SP), when these two sound attributes become correlated, one neuron’s responses would become predictable from the responses of the other, making one neuron’s responses redundant. Efficient coding theory predicts that as stimulus parameters become correlated, tuning functions of different neurons should become less alike (or decorrelated) so as to reduce response correlations (and predictability) and increase coding efficiency (Barlow and Földiák, 1989). To test this conjecture, we calculated tuning functions along AM and SP dimensions separately for each unit. Then, we paired *simultaneously* recorded neurons, and calculated the correlation coefficient between the AM tuning function in one neuron and the SP tuning in the other (**Figure 5A**), both for the Pre-exposure and Post-exposure tests, and finally computed the difference in the correlations before and after exposure. Further, because correlations estimated from different neuron pairs in each recording were not independent, we averaged changes in correlation coefficients from all simultaneously recorded neuron pairs for statistical analysis. We hypothesized that exposure to positively correlated attributes would lead to a decreased correlation coefficient, while exposure to negatively correlated attributes would lead to an increased correlation coefficient, because of de-correlation of initially negatively correlated tuning functions.

The results are plotted in a cumulative frequency distribution (**Figure 5B**). After exposure to positively correlated attributes, the majority of recordings (8/11) exhibited reduced correlations between AM and SP (**Figure 5B**, blue trace), while after exposure to negatively correlated attributes, 7/10 recordings exhibited positive correlation changes (**Figure 5B**, red trace), indicating a significant contrast between the two groups (Mann–Whitney U test, *p* = 0.018). **Figure 5C** summarizes the results in a box plot. Note that the opposite direction of changes after exposure to negatively-correlated attributes and positively-correlated attributes actually reflects the same basic effect: tuning functions along the AM and SP dimensions in different neurons became decorrelated, or more dissimilar, after exposure. As a control, we also computed the correlations between AM-tuning function and S-tuning functions within the same neuron for positively- and negatively-correlated exposures (**Figure 5D**). The results showed no significant difference in the correlations between the two types of exposures (**Figure 5E and 5F**, Mann–Whitney U test, *p* = 0.674).

The neural response changes we describe strongly support Barlow’s hypothesis that neural tuning to correlated properties becomes de-correlated in the population of neurons after sufficient exposure to the covariance, even in the passively listening animal. Neural mechanisms to account for this phenomenon can be readily simulated through a computational model of a population of neurons (N = 200) that are arbitrarily and uniformly tuned to a two-dimensional of stimulus matrix (100 AM steps X 100 SP steps) similar to the test-stimuli matrix in our experiments. Details of a specific implementation are shown in **Figure 5G**. By assuming that the effect of exposure is to move each neuron’s tuning function away from the parameters of the exposure stimulus (red dots in **Figure 5G**) along the orthogonal direction (as indicated by the green and blue arrows), one can recreate the effect of decorrelations (**Figure 5H**) and tuning function shifts (**Figure 5I**).

### Experiment 2: The effects of exposure to stimuli on a single feature dimension

In this experiment, we wanted to determine whether exposure to covariance of two properties was necessary to cause the adaptation patterns described in Experiment 1. Therefore, we recorded from 56 single-units in A1 in two ferrets as they were exposed to stimuli varying along one dimension only, AM or the SP, (**Figure 1E**).

First, we found that some of the adaptation and tuning effects were comparable to those of Experiment 1 in that responses to the exposure stimuli (red dots in **Figure 6A**) were significantly reduced (Wilcoxon test: *z* = −2.3. *p* = 0.022) compared to the increased response to the non-exposed stimuli (blue dots in **Figure 6A**; Wilcoxon test: *z* = − 2.5. *p* = 0.015), as illustrated by the histograms and Box plots in **Figures 6B and 6C**. Thus, exposure to sounds varying along one dimension caused an adaptation effect similar to that in Experiment 1 (**Figure 2B and 2C**).

**Figure 6.**
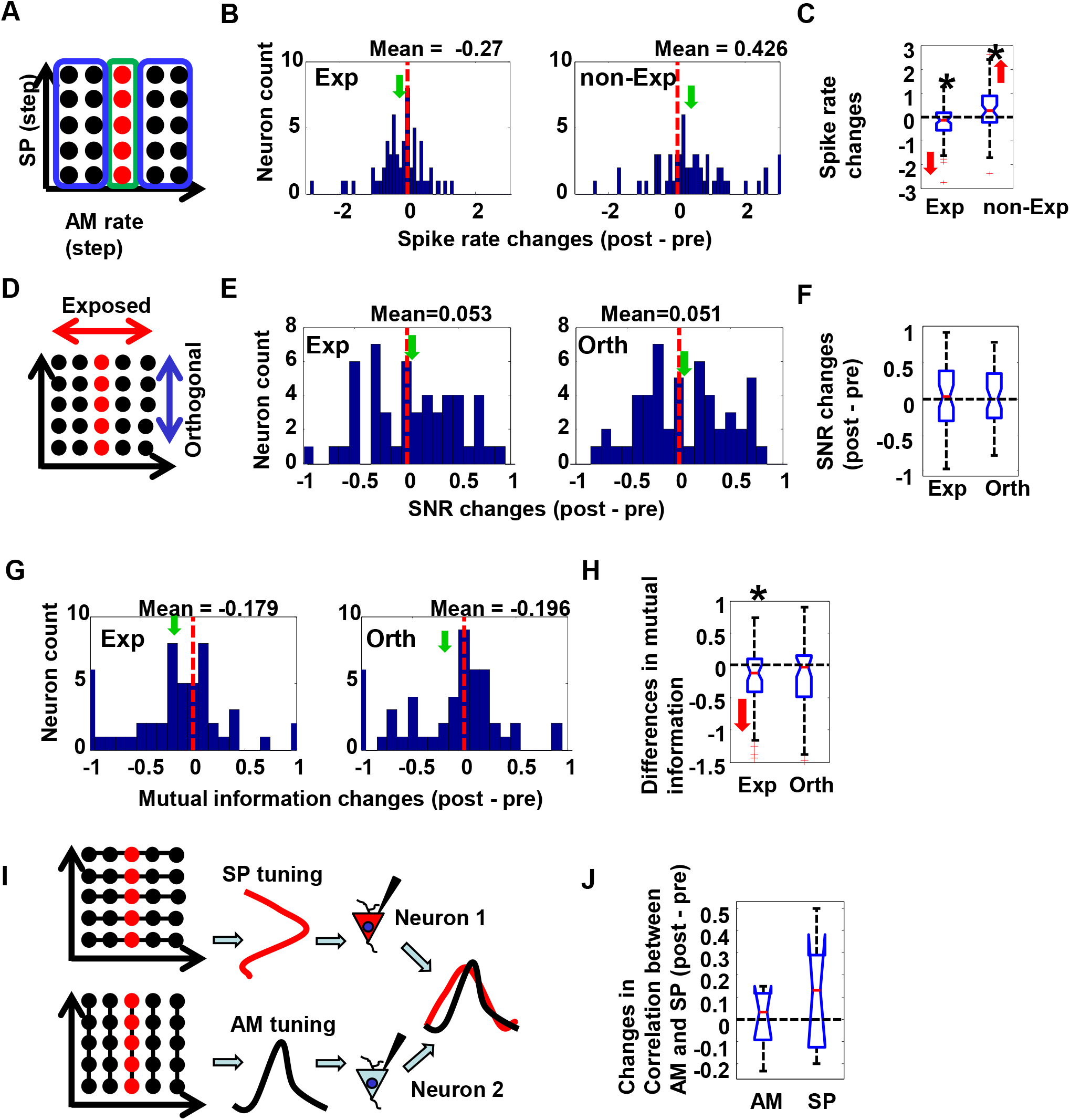
A. Exemplars of stimuli used in tuning analysis in Experiment 2. Green rectangle indicates exposure stimuli, from which the spike rates were averaged and compared between tests before and after exposure. Two blue rectangles indicate non-exposure stimuli, from which the spike rates were averaged for comparison. B. Histogram of spike rate changes after exposure in Experiment 2. Spike rate significantly decreased for the exposure stimuli (Left) and significantly increased for non-exposure stimuli (Right), which resembled results in Experiment 1. C. Box plot of spike rate changes after exposure in Experiment 2. Spike rate significantly decreased for the exposure stimuli (left) and significantly increased for non-exposure stimuli (right). D. Schematic illustration of the stimuli tested in Experiment 2. Red dots indicated exposure stimuli that were only changed in a single dimension (AM or SP, balanced across recordings). Red arrows indicated the exposed dimension and blue arrows indicated the orthogonal dimension. E. Histogram of normalized SNR changes in Experiment 2. Both the distribution for stimuli along exposure dimension (left) and the distribution for stimuli along orthogonal dimension (right) were symmetrically around zero, without any significant bias. The means of the distributions are indicated both on the top of the plot and by the green arrow. F. The box plot of normalized SNR changes in Experiment 2. The median of SNRs for both dimensions overlapped with zero. Thus, there was no change in SNRs in either dimension in Experiment 2. G. Histogram of MI changes in Experiment 2. Contrary to the results of Experiment 1, MI significantly decreased for exposure stimuli in Experiment 2. H. The box plot of MI changes in Experiment 2. I. Schematic illustration of calculation of correlation of tuning from different neurons in Experiment 2. J. The box plot of tuning correlation changes in Experiment 2: There was no significant change in tuning correlation for either of the two types of exposure.

However, unlike Experiment 1, comparing the SNR changes along the exposure vs orthogonal dimensions (**Figure 6D**) revealed *no* significant changes in coding quality, as summarized by the histograms of **Figures 6E and 6F** (Wilcoxon test: *z* = −0.1. *p* = 0.940 for exposure responses, and Wilcoxon test: *z* = −0.25. *p* = 0.800 for the unexposed stimuli). Therefore, it is evident that the SNR effects observed in Experiment 1 were dependent on exposure stimuli with correlated-attributes.

Similarly, we measured effects of single-dimension exposures on MI changes. In this condition, MI on the exposure dimension significantly *decreased* (Wilcoxon test: *z* = −2.4. *p* = 0.016) contrary to the results of Experiment 1, as shown in **Figure 5G and 5H**. There was no significant change in MI on the orthogonal dimension in both Wilcoxon test (*z* = −1.6. *p* = 0.109) and binomial test (binomial test: *p* = 0.200). Considering this difference between the results of Experiment 1 and Experiment 2, we concluded that exposure to correlated stimuli was necessary to elicit changes in MI. Finally, we also analyzed changes in the correlation between the tuning curves (**Figure 6I**). Data from AM and SP exposures were separately analyzed and compared, and no significantly different correlation changes were found between the two types of exposure (**Figure 6J**, Mann–Whitney U test, *p* = 0.268).

In summary, exposure to sounds varying along one dimension in Experiment 2 led to adaptation and tuning shifts as in Experiment 1, but did not lead to significant changes in SNR, MI, or correlations between neurons, as had been observed in Experiment 1.

### Experiment 3: Coding efficiency with exposure to stimuli of correlated features embedded in a higher-dimensional space

Here we extended Experiment 1 to explore whether the same adaptive effects induced by exposure to stimuli of two correlated features would persist if the stimuli had an additional, but uncorrelated, third acoustic feature (e.g., *f*0) that varied randomly.

Responses were recorded from 56 A1 single-units in two ferrets that were exposed to stimuli located at the two diagonals of the testing space (**Figure 7A**), axes along which we previously found the strongest adaptive effects. The results obtained strongly resembled those of Experiment 1: (1) SNRs decreased for the orthogonal dimension (**Figure 7B and 7C**, Wilcoxon test: *z* = −2.3. *p* = 0.02), while remaining intact for the exposure dimension (Wilcoxon test: *z* = −0.4. *p* = 0.677). (2) MI’s increased for the exposure dimension (**Figure 7D and 7E**, Wilcoxon test: *z* = −2.1. *p* = 0.035; binomial test: *p* = 0.041), while remaining unchanged for the orthogonal dimension (Wilcoxon test: *z* = −1.2. *p* = 0.250; binomial test: *p* = 0.17). When data from Experiments 1 and 3 were combined, MI’s increased for the exposure dimension (Wilcoxon test: *z* = −2.7. *p* = 0.008; binomial test: *p* = 0.003) but remained unchanged for the orthogonal dimension (Wilcoxon test: *z* = −1.0. *p* = 0.299; binomial test: *p* = 0.293).

**Figure 7.**
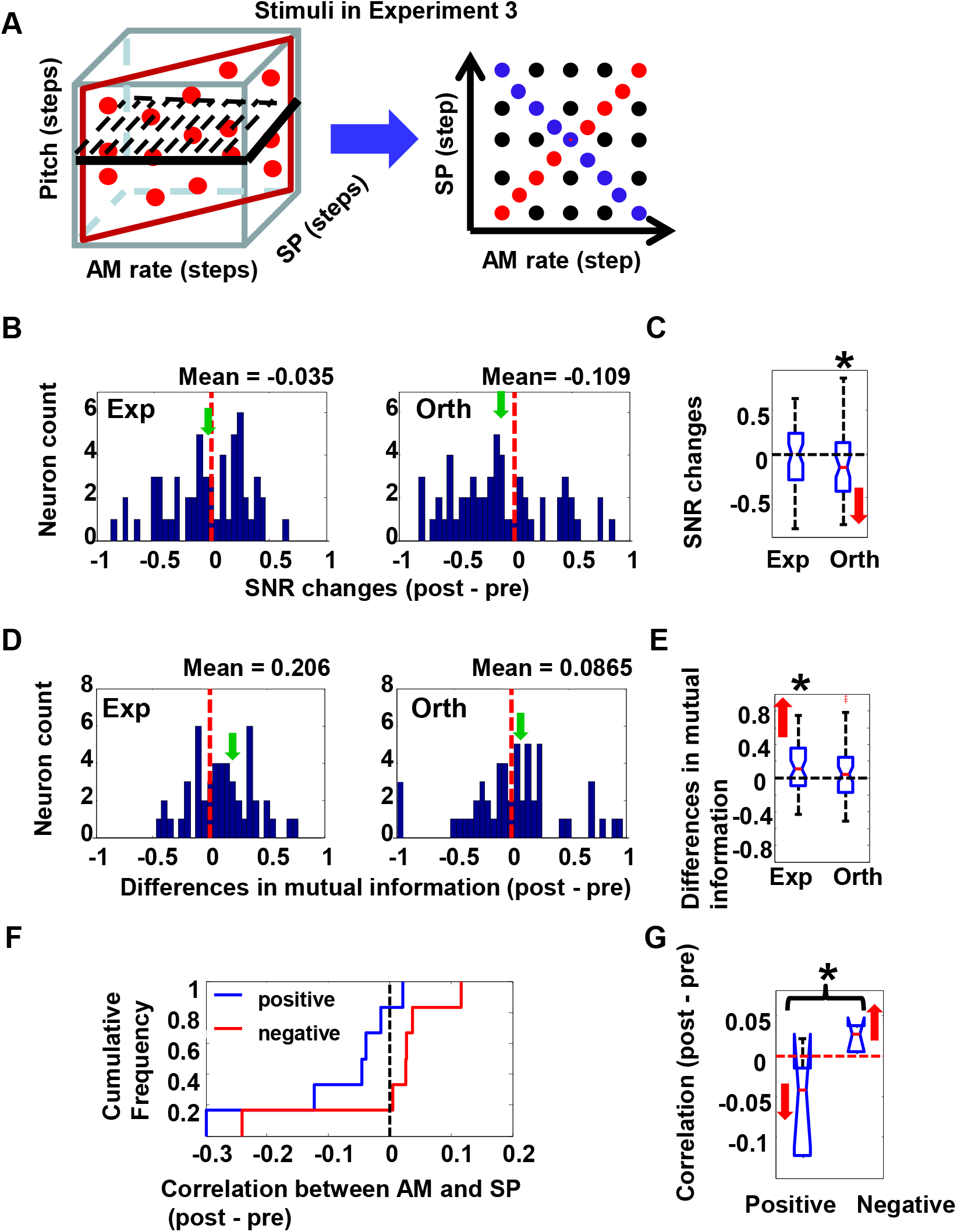
A. Testing stimuli space in Experiment 3. The red parallelogram indicates exposure sounds, which had perfect positive or negative correlation between AM and SP, and varied f0s. Black parallelogram with dashed lines indicates the testing sounds, in which only one pitch was randomly chosen for testing in each recording. The structure of the testing space was same as in Experiment 1, indicated on the right by the blue arrow. B. Histogram of SNR changes in Experiment 3. SNRs for exposure dimension were symmetrically distributed around zero (left, red dashed line indicates zero). Consistent with Experiment 1, SNRs for orthogonal dimension were significantly reduced (right), resembling what was found in Experiment 1. C. Box plot of normalized SNR changes in Experiment 3. D. Histogram of MI changes in Experiment 3. Consistent with Experiment 1, MI significantly increased for exposure stimuli (left), but not for orthogonal stimuli (right). E. Box plot of MI changes in Experiment 3. F. Cumulative frequency distribution of tuning correlation changes across neurons in Experiment 3: Consistent with Experiment 1, the blue trace showed that correlation decreased (5 out of 6) after exposure to positively correlated attributes. The red trace showed that correlation increased (5 out of 6) after exposure to negatively correlated attributes. The contrast between the two distributions was statistically significant. G. Box plot of MI changes tuning correlation changes across neurons in Experiment 3.

Finally, in Experiment 3, correlation of tuning functions to AM and SP between neurons decreased in 5/6 recordings following exposure to positive covariance and increased (in 5/6 recordings) after exposure to negative covariance (**Figure 7F and Figure 7G**, Mann–Whitney U test, *p* = 0.032). Combining data from Experiments 1 and 3, correlations decreased in 13/17 recordings after exposure to positive covariance (binomial test: *p* = 0.038), while 12/16 recordings showed increased correlation after exposure to negative covariance (binomial test: *p* = 0.025). Therefore, exposure to sounds with an additional randomly varying property like *f*0 did *not* affect the efficient coding of the correlated features, consistent with the results in Experiment 1.

## Discussion

Experiments described here sought to explore the neural correlates underlying the phenomenon of adaptive coding of contingent sensory properties, which was proposed as a key principle of coding in sensory systems. In vision, this principle is exemplified by the McCollough effect in which the visual system, following passive exposure to correlated properties of visual stimuli (color and orientation), combines them as a single property (McCollough, 1965). The physical acoustics of natural sounds often reveal correlations between acoustic attributes (Lutfi et al., 2011). In auditory perception, the adaptive coding principle is also supported by the results of a set of psychoacoustic experiments that were the inspiration for this study (Stilp et al., 2010; Stilp and Kluender, 2011, 2012, 2016).

The adaptive effects we observed in the neural recordings in the auditory cortex of the ferret were consistent with, and mapped readily to the results of the psychoacoustic experiments in human subjects. Furthermore, the changes allowed us to explore the underlying mechanisms that gave rise to this coding efficiency, and whether this neural adaptation was resilient to the addition of other uncorrelated feature dimensions. Our findings indicate that two aspects of the adapted responses contribute to the significant increase in coding efficiency after exposure to stimuli with the AM and SP correlated properties: (1) in single neurons, repeated exposure to stimuli of correlated attributes reduced their responses with a concomitant decrease in SNR of responses to stimuli along the orthogonal dimension. This change was combined with an enhanced MI between spike trains and stimuli, such that fewer spikes conveyed information about the correlated stimuli efficiently in their trains. (2) Tuning curves of different neurons to the attributes of the AM and SP stimuli became less correlated following exposure, enhancing the efficiency of the population coding of stimulus identity. Consequently, as seen in Experiment 2, the exposure effects did not materialize when stimulus parameters varied along only one dimension (AM or SP); although, adaptation of spike rates still occurred to the exposure stimuli as before. Finally, we found in Experiment 3 that neurons were capable of extracting and enhancing the coding efficiency of two correlated properties even when they were embedded in a higher dimensional space that included independent variation in a third dimension.

Reduced SNR along the orthogonal dimension reflects an adaptive coding mechanism in single neurons in response to dynamically changed feature statistics. This mechanism can be modeled by changes in the receptive field (RF) of single neurons which are approximated by 2-D Gaussian probability density function (PDF) as shown in **Figure 3c**. The variance of the PDFs is estimated by the mean of the variances between stimuli along the exposure and orthogonal dimensions, which were calculated from the 65 neurons in Experiment 1. Because there were no significant changes in overall response amplitude, changes in PDF widths before and after exposure illustrate how such change may contribute to adaptive coding of a new combined feature that emerged during passive exposure. As shown in **Figure 8**, the estimated RFs after exposure became tilted along the exposed dimension (middle). To examine this change further, we took the difference between the two PDFs (panel on the right). The RF change reflects increased responsiveness that takes the shape of an ellipse along the exposure dimension, with reduced responsiveness along the orthogonal dimension. Such a change would optimize a neuron’s capacity to capture the emergent feature along the exposure dimension at the cost of coding capacity along the orthogonal dimension. This is analogous to principle component analysis (PCA), in which the coordinate of variance measurement is altered to capture the most important feature. This hypothesis is consistent with behavioral results and modeling by Stilp et al., (2010) and Stilp and Kluender (2012, 2016). Notice that, as the RF shapes change, the tuning center of each neuron also moves away from the exposed diagonal in the testing feature space (as was modeled in **Figure 5G**), which in turn leads to an overall decrease in response amplitude for exposed stimuli (**Figure 2D**).

Previous studies have shown that sensory systems can combine different sensory cues to improve perception. For example, in vision, shape information from texture, and depth information from disparity cues, can be combined to form a fused percept, leading to improved discrimination performance compared to performance based on single-cues (Hillis et al., 2002). In haptic perception, force and position cues can be integrated for shape perception (Robles-De-La-Torre and Hayward, 2001; Drewing and Ernst, 2005). Furthermore, sensory cue integration may even occur across different sensory modalities (Ernst, et al. 2000; McGurk and MacDonald, 1976; Shams, et al. 2000).

We should note that the adaptive effects we have studied here are induced by short periods of passive exposure (although the effects may become enhanced with behavioral training and/or task engagement). Thus, the ability to capture regularities though repetitive passive exposure may not only benefit the organism in sensory coding efficiency, but also function as a simple form of learning. There is considerable evidence that sensory systems can learn different types of regularities through such passive exposure: transition probability in sound sequences (Saffran et al., 1996, Newport and Aslin, 2004; Lu and Vciario., 2014) and other complex patterns (Agus et al., 2010; McDermott et al., 2011; Barascud et al., 2016; Lu et al, 2018). Our current study fits into this mold, and demonstrates the underlying neural transformations that make it possible. We note that our observations in this study are also interpretable in the context of implicit learning and habituation (Lu et al, 2018) in which exposed stimuli are contrasted with novel (unexposed or “orthogonal”) stimuli. However, thoroughly exploring this interpretation of our results will require a new study in which exposures and analyses are adapted to the parameters typical of statistical learning paradigms (Lu et al, 2018).

Pre-verbal infants, nonhuman primates and birds can learn and recall different artificial grammars and show implicit long-term statistical learning of auditory patterns and sequences (Saffran et al., 1996; 1999; Hauser et al.,2001; Newport et al., 2004; Abe et a., 2011). For example, monkeys implicitly learn acoustic presentations of simple artificial grammars after an exposure of 20-30 minutes (Fitch and Hauser, 2004; Wilson et al., 2013). Human adults show an extraordinarily robust and long-lasting auditory memory that can develop rapidly for random noise or time patterns (Agus et al. 2010; Kang et al. 2017). Human adults can also recover sound patterns from embedded repetitions that were masked by noise (McDermott et al., 2011, Stilp et al., 2018). Human speech is a complex signal characterized by many correlated attributes arising from the physical acoustic constraints of the vocal apparatus and the dynamics of articulating sequences of sounds (leading to co-articulation correlated context effects). Thus it is likely that these multiple correlations (such as power spectrum and manner of articulation (Llanos et al., 2017)) play an important role in the acquisition and the neural representation of human speech.

Our study adds to previous research exploring the brain areas and neural mechanisms underlying implicit statistical learning. Single-unit studies showed that the probability of occurrence of an oddball tone in the context of standard tones is encoded in stimulus-specific adaptation in A1 (Ulanovsky et al., 2003; 2004; Yaron et al., 2012; Nieto-Diego and Malmierca, 2016; Parras et al. 2017). Transition probability in sound sequences is encoded in higher auditory forebrain of songbirds (Lu and Vicario, 2014). Studies of mismatch negativity (MMN) have demonstrated that the brain can detect the violation of acoustic sequences, even when the embedded patterns were very complex (Paavilainen et al., 2007, Barascud et al., 2014). Detection of acoustic regularities activates a network including A1, and higher auditory cortex, hippocampus, and inferior frontal gyrus (Barascud et a., 2016). We note that implicit learning is widespread, and is not limited to the auditory system. For example, human subjects can implicit learn visual transition probability in visual scene sequences. Neuroimaging revealed activation of striatum and medial temporal lobe to learned statistical structure (Turk-Browne et al., 2009). Mice can learn spatiotemporal sequences through implicit learning at the neuronal level in V1 (Gavornik and Bear, 2014). Our current work demonstrates a form of implicit learning, at the neuronal level in A1, that binds features from two correlated attributes, even when the pattern was embedded in high dimensional variance. It is highly likely a larger network, including higher auditory areas, also contribute, and a future challenge is to unravel the roles of different brain areas to implicit statistical learning.

**Figure 8.**
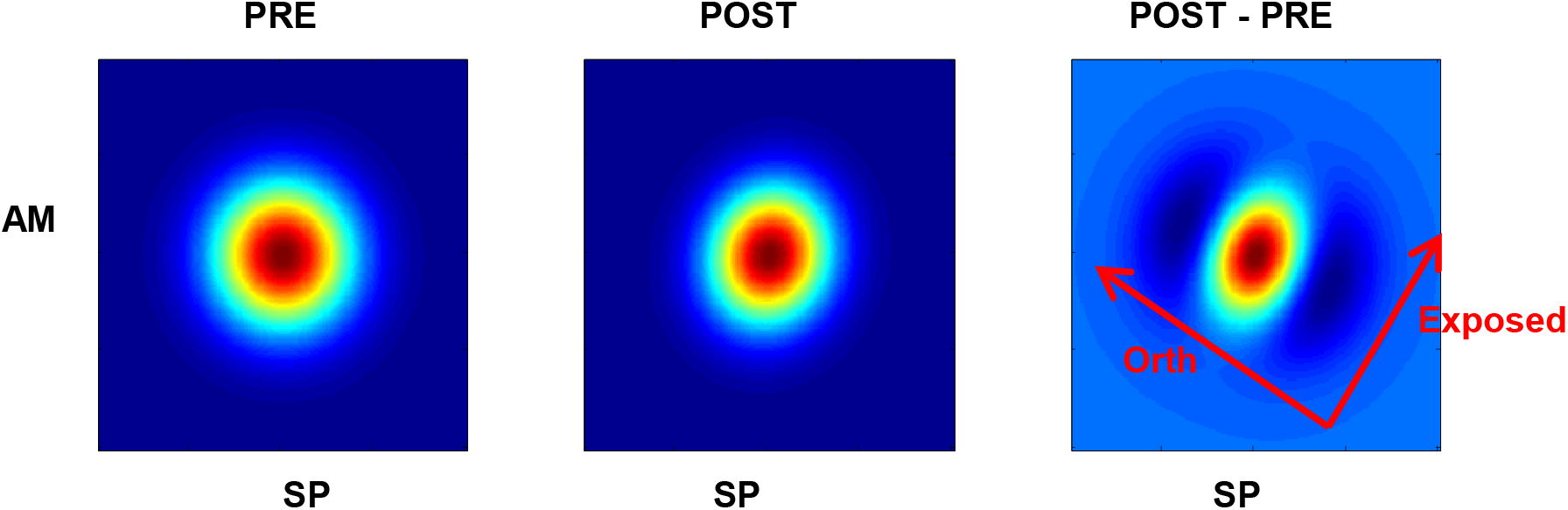
A. Receptive field of neurons before exposure, modeled with 2-D Gaussian probability density function. Variance between stimuli on the exposed dimension and the orthogonal dimension in Experiment 1 were used in this model. B. Receptive field of neurons after exposure, modeled with variance between stimuli from Experiment 1. C. Changes of Receptive fields, by taking the difference between the probability density function before and after exposure. The coordinate of the exposed and orthogonal dimension is indicated by red arrows.

## Acknowledgements

We appreciate the insightful and detailed comments from Dr. Keith Kluender. This research was funded by grants from the National Institutes of Health (R01 DC005779) and an Advanced ERC grant (NEUME) to SAS

